# Differential expression analysis of RNA sequencing data by incorporating non-exonic mapped reads

**DOI:** 10.1101/016196

**Authors:** Hung-I Harry Chen, Yuanhang Liu, Yi Zou, Zhao Lai, Devanand Sarkar, Yufei Huang, Yidong Chen

## Abstract

**Background:** RNA sequencing (RNA-seq) is a powerful tool for genome-wide expression profiling of biological samples with the advantage of high-throughput and high resolution. There are many existing algorithms nowadays for quantifying expression levels and detecting differential gene expression, but none of them takes the misaligned reads that are mapped to non-exonic regions into account. We developed a novel algorithm, XBSeq, where a statistical model was established based on the assumption that observed signals are the convolution of true expression signals and sequencing noises. The mapped reads in non-exonic regions are considered as sequencing noises, which follows a Poisson distribution. Given measureable observed and noise signals from RNA-seq data, true expression signals, assuming governed by the negative binomial distribution, can be delineated and thus the accurate detection of differential expressed genes.

**Results:** We implemented our novel XBSeq algorithm and evaluated it by using a set of simulated expression datasets under different conditions, using a combination of negative binomial and Poisson distributions with parameters derived from real RNA-seq data. We compared the performance of our method with other commonly used differential expression analysis algorithms. We also evaluated the changes in true and false positive rates with variations in biological replicates, differential fold changes, and expression levels in non-exonic regions. We also tested the algorithm on a set of real RNA-seq data where the common and different detection results from different algorithms were reported.

**Conclusions:** In this paper, we proposed a novel XBSeq, a differential expression analysis algorithm for RNA-seq data that takes non-exonic mapped reads into consideration. When background noise is at baseline level, the performance of XBSeq and DESeq are mostly equivalent. However, our method surpasses DESeq and other algorithms with the increase of non-exonic mapped reads. Only in very low read count condition XBSeq had a slightly higher false discovery rate, which may be improved by adjusting the background noise effect in this situation. Taken together, by considering non-exonic mapped reads, XBSeq can provide accurate expression measurement and thus detect differential expressed genes even in noisy conditions.

## Background

Next-generation sequencing (NGS) has been widely used in biological studies. RNA sequencing (RNA-seq) is the most commonly used NGS technologies to investigate the aberration of mRNA expression in disease and normal condition comparison. Unlike microarray technology, which uses a short section of a gene as a probe to determine the gene’s expression, RNA-seq provides measurement across entire exonic region, enabling accurate expression quantification and discovery of novel isoforms and splicing junctions. With RNA-seq technology, thousands of novel coding and non-coding genes, alternative splice forms of known genes have been discovered.

Differential expression (DE) analysis using RNA-seq is commonly employed to interrogate changes between different experimental conditions. While enormously successful, DE analysis also suffers from systematic noise and sequencing biases, such as sequence quality bias, wrong base calls, variability in sequence depth across the transcriptome, and the coverage depth differences of replicate samples [1]. There are already many statistical testing methods for RNA-seq differential expression analysis. One is to normalize the read counts of target transcripts, converting them into reads per kilobase per million mapped reads (RPKM) and then perform linear modeling methods that are used in microarray experiments [2]. However, the method designed for microarray measurement may not fit the characteristics of sequencing data properly. In past years, algorithms have been developed specifically for RNA-seq data analysis. Among them, two popular software packages implemented the negative binomial (NB) model that account for genome-wide read counts and moderate dispersion estimates with different statistical methods [3, 4]. EdgeR [4] uses a trended-by-mean estimate to moderate dispersion estimates, whereas DESeq [3] takes account of the maximum of a fitted dispersion mean or the feature-wise dispersion estimate, as reviewed in [5]. However, neither of these methods considered the misaligned reads existing in the sequencing data, which may play a significant role in detecting the significance of target transcripts.

Here we propose a novel DE analysis algorithm – XBSeq, which is derived from DESeq, where we take the non-exonic reads of RNA-seq data into consideration. In conventional RNA-seq analysis, reads mapped to the exons are counted as the expression of a gene, whereas reads aligned to the intronic and inter-genic regions are generally ignored. Those non-exonic hits exist because of: sequencing error, mapping error, contamination by genomic DNA, unannotated genes, and nascent transcription and co-transcriptional splicing [6]. Our model treats these sequence reads as sequencing noises that exist across the entire genome, both exonic and non-exonic regions. Therefore, the observed read counts can be decomposed into two components: true signals that are directly derived from transcripts expression, and the others from the random noises. We model true expression signals by a negative binomial distribution and assume sequencing background noises possess a Poisson distribution. With non-exonic read counts, we can estimate the parameter *λ*_*i*_ of the Poisson distribution of each gene. Afterwards, we remove sequencing noise effect from observed signals and retrieve the background-corrected mean and variance parameters for the NB model of true expression signals.

To study the robustness of the algorithm, we have built a simulation framework that generates RNA-seq data by combining the true signals in NB distribution with different levels of non-exonic reads from Poisson distribution. We demonstrate our method by applying it to the simulated data and examine how it performs comparing with other common DE analysis algorithms.

## Methods

### Non-exonic sequence read count

For a typical transcriptome profiling by RNA-seq, we detect read count of each gene by using HTSeq algorithm [7]. Given exons’ locations of every gene, HTSeq counts sequence reads aligned to the genic regions. In order to capture the reads in non-exonic regions, we preserved the structure information of each gene (transcript length, exon size, etc) by shifting start and end positions of each exon to a nearby intronic or inter-genic region (See Fig. 1A). We have defined non-exonic regions for each species by the following steps:

**Figure 1.**
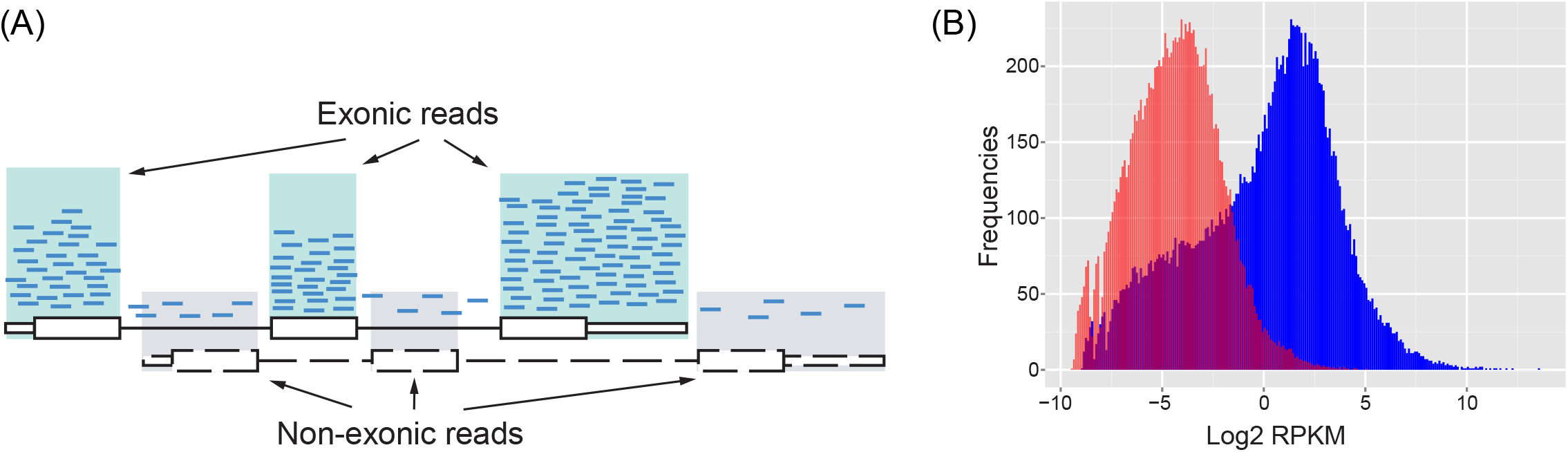
(A) Illustration of exonic and non-exonic reads. (B) Histogram of sequence read counts in RPKM. The histogram of observed signal (*X*) is plotted in blue and the histogram of non-exonic read counts (*B*) in pink.

1. Download refFlat table from UCSC database (https://genome.ucsc.edu) and create the preliminary list of gene-free regions,
2. Download tables of (a) all_mrna; (b) ensGene; (c) pseudoYale60Gene; (d) vegaGene;, (e)xenoMrna, and (f) xenoRefGene from UCSC database and remove regions appear in any of them from the gene-free regions,
3. To guarantee gene-free regions are far enough from exonic regions, trim 100 bps from both sides of intronic regions and 1,000 bps from both sides of inter-genic regions,
4. Shift each exon of a gene to the right nearest gene-free region. Most of the shifted genes remain the same as the original structures of the genes,
5. If the nearby gene-free region is too short, we may only preserve the exon size features but not the whole gene structure. The priority of shifting a region is: i) nearest right gene-free region, 2) nearest left gene-free region; 3) the second right nearest gene-free region and so on until the shift region of the original exon fits, and
6. At last, we considered the shifted regions as the non-exonic regions for each gene and a final .gtf file was created.

To extract non-exonic read counts, HTSeq was performed second time to generate an equivalent read count for each gene over an exactly same length of non-exonic region. By doing so we guarantee an equivalent read count from non-exonic region for each gene.

The histogram of a RNA-seq data in RPKM unit was plotted in Fig. 1B. The blue histogram was derived from the observed read counts genome-wide, while the red one was derived from the non-exonic read counts after shifting the exons’ position. As illustrated in the figure, the hump of the red histogram overlaps with the left tail of the blue histogram, indicating the existence of sequencing noises in commonly reported gene expression levels, particularly when gene expression level is low. Based on this observation, we hypothesize that the read count of *i*^th^ gene (e.g., observed by HTSeq), defined as *X*_*i*_, is composed of true signal *S*_*i*_ (not measurable directly) and background noise *B*_*i*_ (measured over our uniquely defined non-exonic regions).

### Poisson-Negative Binomial model

We assume that read count of *i*^th^ gene can be decomposed into two components: true signal *S*_*i*_ that directly derived from transcript expression, and background noise *B*_*i*_ due to sequencing error or misalignment. Therefore, the observed signal *X*_*i*_ is

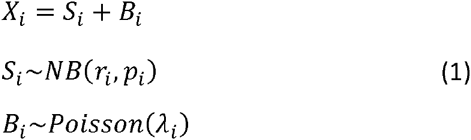

where *r*_*i*_, *p*_*i*_ are parameters (number successes and probability of success, respectively) of NB distribution and *λ*_*i*_ is the rate parameter of Poisson distribution. We further assume that the true expression signal *S*_*i*_ and background noise *B* are independent. Given the observed signal *X*_*i*_ to be the sum of a NB and a Poisson, the probability distribution of *X*_*i*_ is governed by a Delaporte distribution, which is the convolution of a NB distribution with a Poisson distribution [8, 9]. When there is no background noise (which is the assumption of many other RNA-seq algorithms), the observed signal is simply governed by a NB distribution,

### Estimation of distribution parameters

The two NB parameters *r*_*i*_ and *p*_*i*_ can be estimated by the background corrected mean and variance of gene *i*; and with the non-exonic read counts, the Poisson parameter *λ*_*i*_ can be determined easily. We further assume that genes are independent to each other, acknowledging that some genes are dependent within pathways or other reasons. Hence, the objective is to estimate all parameters of each gene in order to obtain the NB model fitted to the true expression signals.

The estimation of Poisson parameter *λ* is relative simple, we assume that the read count derived from the non-exonic regions representing the background component *B* and independent of *S.* Therefore, we can obtain *λ* of each gene, being the average of total non-exonic read counts across all *m* replicates.

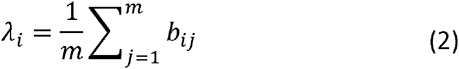

where *b*_*ij*_ is the non-exonic read count of *i*^th^ gene from *j*^th^ sample. After the estimation of Poisson parameter, we can calculate the true expression signals’ mean *μ*_*Si*_ and variance *σ*_*Si*_ of each gene as follows,

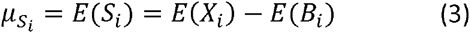

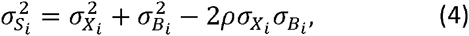

where 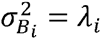. Note that observable *X* and background *B* are not independent. As we mentioned earlier, observed read count *X* follows a Delaporte distribution, which has no closed form [8, 9]. The parameters of Delaporte distribution, D(λ, α, β), however, is known as *μ* = *λ* + *αβ*, and *σ*^2^ = *λ* + *αβ*(1 + *β*), where *λ* can be considered as the parameter of Poisson distribution. When *λ* = 0, the Delaporte reduces to NB distribution, similar to what we have in Eqs. (3) and (4). The same variance correction method in DESeq is subsequently used to adjust 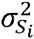 in order to get precise estimate of the variance when the number of replicates is small [3]. After obtaining the adjusted true expression signal mean and variance, *μ*_*S*_ and 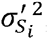, from Eqs. (3) and (4), the two NB parameters *r*_*i*_ and *p*_*i*_ are further estimated by,

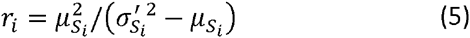

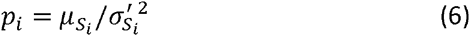

### Testing for differential expression

After estimating the NB parameters in both experimental conditions, differential expression analysis between the two conditions can be tested. Designed similarly to DESeq method, we use a Fisher’s exact test approach to estimate the *P* value of each gene [3]. In short, suppose we have *x* and *y* reads of a gene in each condition, we compute every possible *p(a, b)*, where the sum of the variables *a* and *b* equal to *K*_*total*_ (*K_total_ = x + y*). By assuming the independence of two test conditions, we have *p(a, b) = Pr(a) Pr(b)*, where *Pr(a)* and *Pr(b)* are the probabilities in NB distribution that we have estimated for each condition. Therefore, the *P* value is the proportion of the sum of possible probabilities less than the probability of actual read counts among the sum of all probabilities as follows,

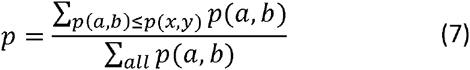

Equation 7 is evaluated gene-wise, and for simplicity, we omitted subscript *i*.

### Simulation

In order to evaluate the performance of different RNA-seq algorithms, we generated a set of simulated data where we could control the differential expression status for a given set of genes, as well as noise level for all genes. In this study, true signal *S* and background signal *B* were simulated based on a negative binomial distribution and a Poisson distribution, respectively, with parameters estimated from real RNA-seq data.

We followed a similar simulation framework used by edgeR-robust [5]. Firstly, genes from real RNA-seq data were filtered based on the expression intensity across all replicates. The genes with top 10% dispersion were discarded. Then 5,000 genes were randomly selected with replacement among the filtered genes. Based on the mean and dispersion estimated from real RNA-seq data, the true signal *S was* simulated for each gene from the negative binomial distribution. Different proportions of genes (10% and 30%) were randomly selected as differentially expressed genes with various fold changes (1.5, 2, 3, and 5). To simulate baseline background signal *B*_*baseline*_, firstly, the mean value was calculated using the non-exonic mapped reads of its corresponding gene. Then *B*_*baseline*_ was generated from a Poisson distribution with parameter *λ* equals to the calculated mean value from the non-exonic read counts. Observed signal *X*_*baseline*_ was then generated as the addition of *S* and *B*_*baseline*_.

To simulate background signal *B*_*inc*_ with increased non-exonic mapped reads, we first calculated mean read count for each selected gene based on the non-exonic mapped reads from real RNA-seq data. Then *B*_*inc*_ was simulated by a hybrid model,

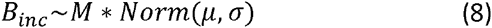

where *μ* is from a Poisson distribution *μ~;Poisson*{*λ* + *NF*), *NF* is the noise factor, which we set to be 0 (low), 7 (intermediate) and 20 (high). *λ* equals to the mean of the non-exonic mapped reads of a given gene, and we set *σ* = 3 and multiply *M* = 10 for our simulation. Finally, we set the observed signal *X*_*inc*_ to be the addition of *S* and *B*_*inc*_. For noise models different from Poisson, we simply replaced Poisson with binomial, uniform or other distributions in Eq. 8. Simulations were performed 100 times in order to evaluate and plot the Receiver Operating Characteristic (ROC) curves and other statistics.

### RNA-seq data set for testing

A mouse RNA-seq dataset were obtained from Gene Expression Omnibus (GSE61875) [10]. For this testing purpose, we selected 3 replicates of wild type mouse liver tissues (WT) and 3 replicates of Myc transgenic mouse liver tissues (MYC) for differential expression analysis to determine differential expressed genes due to the activation of Myc. Out of the six samples selected, on average, 12,781 (out of 22,609) genes have at least one non-exonic reads, and 15,973 genes have at least one exonic reads.

### Comparison of other RNA-seq algorithms

We also compared XBSeq with other differential expression analysis methods, including DESeq (1.14.0) [3], latest version of DESeq (DESeq2, 1.2.10) [11], edgeR(3.5.15) [4], latest version of edgeR (edgeR-robust 3.5.15) [5], baySeq (1.16.0) [12], limma (3.18.13) [13], and EBSeq (1.2.0) [14]. All these evaluation were performed under R version 3.0.2 and Bioconductor version 2.13. Detailed workflow of XBSeq and simulation is illustrated in Fig. 2.

**Figure 2.**
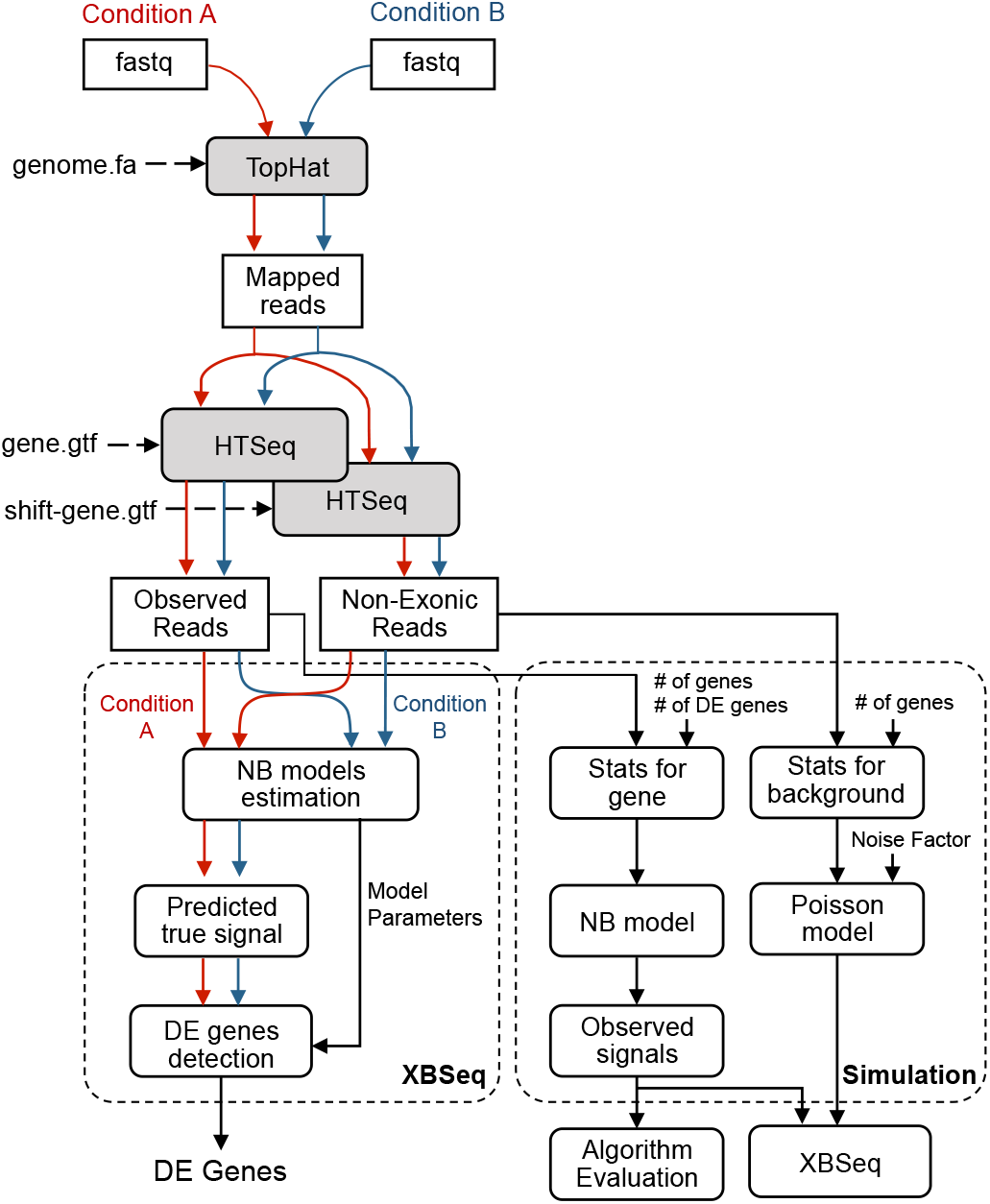
Block diagram of the XBSeq and simulation process.

## Results

### Implementation of XBSeq algorithm

XBSeq requires two inputs, the observed measurement from exonic regions and the background noise from non-exonic regions. Both read counts can be obtained by submitting mapped sequence reads to HTSeq twice with coding gene annotation (*e.g.*, gene.gtf) and shifted gene annotation (shift-gene.gtf) as discussed in Methods Section (Fig. 2). As the first step, the true signal for each gene is estimated by using Eq. 3. The read counts of genes with negative true signals will be automatically assigned to 0. We apply a similar method as DESeq to normalize the true signal based on size factors calculated from each sample, and we then estimate the variance of the true signal for each gene by using Eq. 4. A similar framework for variance correction and the differential expression significance testing as DESeq (Eqs. 5-7) is applied to generate *p*-values for each gene. The output of XBSeq contains *p* value, adjusted *p* value for multiple test correction, log_2_ fold change and other statistical merits for each gene.

To optimize the performance of XBSeq, we applied variance correction procedure either at observed signal level or at estimated true signal level. After examining the performance of these two choices based on simulated datasets, we concluded that variance correction at estimated true signal level yields the best performance with larger area under the ROC curve. We also investigated the performance of LOWESS as well as locfit R package for fitting variance-mean relationship. Locfit was proven to generate more robust variance-mean relationship based on simulated datasets. Therefore, we selected locfit package to estimate variance-mean relationship and carry out variance correction procedure at estimated true signal level in the current XBSeq implementation.

### Discrimination between DE and non-DE genes

To compare the performance of XBSeq with other statistical methods, including DESeq, we generated synthetic data where the variability and fold change of differential expression genes could be controlled. To simulate RNA-seq data with baseline background noise, we generated signal and background read counts (non-exonic region mapped reads) with distribution parameters (negative binomial distribution for true expression signal and a Poisson distribution for background noise) estimated from a set of real RNA-seq data. After removing not expressed genes, we randomly selected 5,000 genes and the number of differentially expressed (DE) genes were set to be 500, 1500, with 1.5 fold, 2 fold, 3 fold or 5 fold changes. To compare XBSeq and DESeq under circumstances with increased non-exonic mapped reads, we carried out the simulation to generate low, intermediate, and high levels of non-exonic mapped reads (see Methods section for detailed simulation procedure and parameter settings). We select DESeq algorithm to compare due to the similarity of statistical evaluation of differential expression levels.

The ability to discriminate between DE and non-DE genes was evaluated by area under the Receiver Operating Characteristic (ROC) curve (AUC). As shown in Fig. 3a and Table 1, when the non-exonic mapped reads are at baseline level, the performance of XBSeq is indistinguishable with DESeq in terms of AUC. Specifically, when the fold change was set to 1.5 with 500 DE genes, both XBSeq and DESeq have equivalent performance with either 3 (AUC = 0.90 and 0.90, respectively) or 6 replicates (AUC = 0.96 and 0.97, respectively). As shown in Table 1, the number of differentially expressed genes had little effect over the performance of XBSeq and DESeq, and both XBSeq and DESeq performed equivalently at different fold changes. When fold change is set to 3 or 5, both methods had no difficulty in detecting virtually all DE genes, with AUC almost equals to 1 (Supplementary Table S2). Overall, under baseline level, the performance of XBSeq and DESeq is comparable.

**Figure 3.**
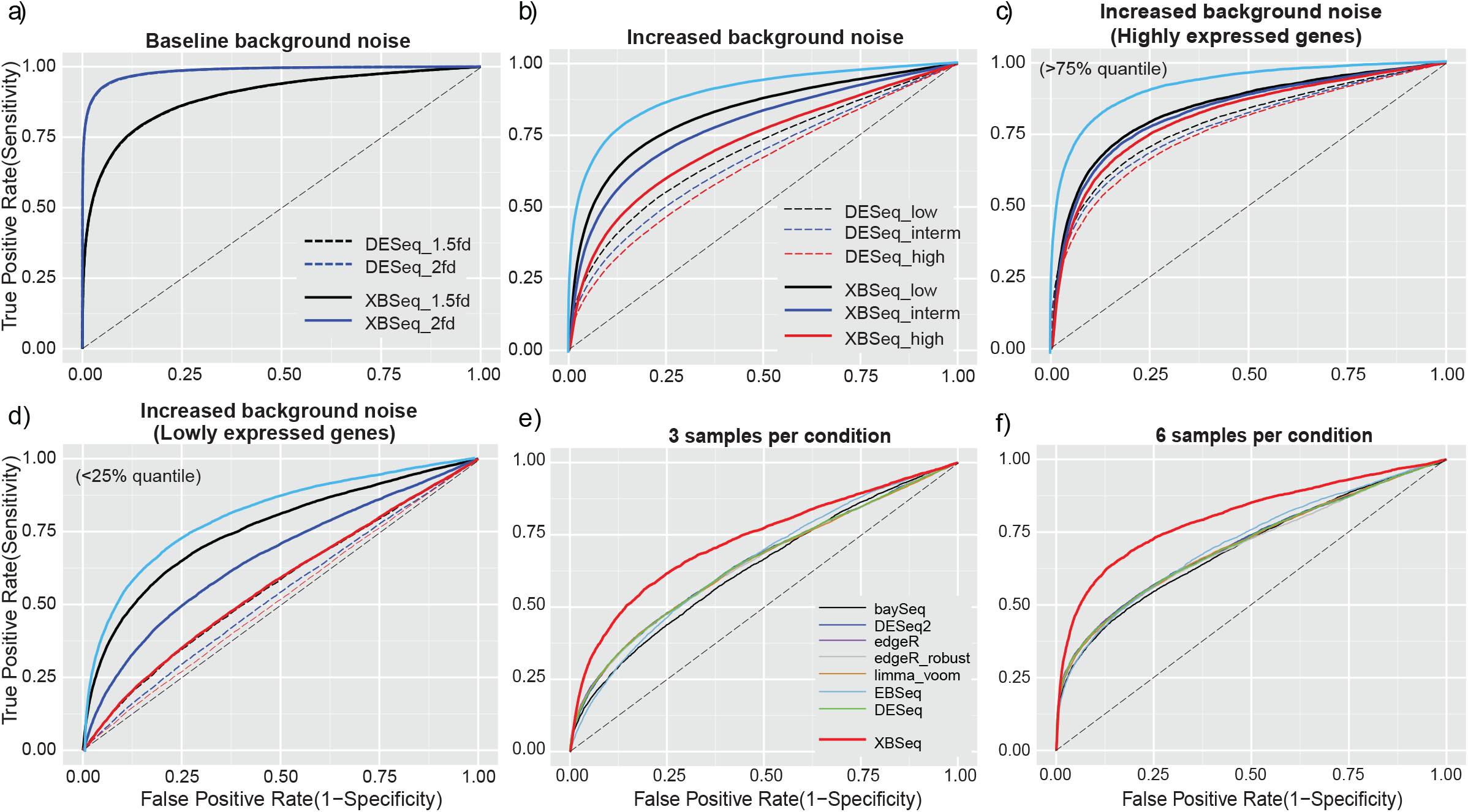
ROC curves of simulated RNA-seq data in different scenarios. ROC curves in the scenarios of baseline background noise (a); Increased background noise (b); Increased background noise but only with highly expressed genes above 75% quantile of intensity (c); Increased background noise but only with lowly expressed genes below 25% quantile of intensity (d); Comparison with other differential expression analysis methods (under increased background noise) with either 3 samples per condition (e), or 6 samples per condition (f). The diagonal black dash-line indicates performance under random events (0.5 AUC). The light blue line in (b,c,d) indicates the theoretically optimal ROC when the background noise can be estimated exactly. Simulation for (a, b, c and d) was carried out 100 times. Simulation for (e and f) was carried out 10 times. 3 replicates per test condition, with 10% DEGs and 1.5 fold change were used for (a, b, c, and d).

**Table 1.** Area under the ROC curve (AUC) for DESeq and XBSeq under various conditions with different number of replicates (3 and 6 replicates), different number of differential expressed genes (500, 1500) and different level of fold change (1.5 fold and 3 fold). Fold change at 5 or higher yield AUC = 1.00 for all conditions.

| Replicates<br>per group | DEGs (%) | DESeq |  | XBSeq |  |
| --- | --- | --- | --- | --- | --- |
|  |  | Fold Change |  | Fold Change |  |
|  |  | 1.5x | 3x | 1.5x | 3x |
| 3 | 10% | 0.90 | 1.00 | 0.90 | 1.00 |
|  | 30% | 0.89 | 1.00 | 0.89 | 1.00 |
| 6 | 10% | 0.96 | 1.00 | 0.97 | 1.00 |
|  | 30% | 0.96 | 1.00 | 0.96 | 1.00 |

With increased background noise, as shown Table 2, as we expected, more replicates per test group performed better in terms of AUC, but the performance decreased as with the increase of the background noise level, but overall, XBSeq has a larger AUC comparing with DESeq in various conditions (AUC_XBSeq_ – AUC_DESeq_ ~ 0.11, on average. See Table 2). For instance, with 6 replicates per test condition, XBSeq achieved AUC of 0.89 with lower background (lower non-exonic read count), while AUC_DESeq_ was only 0.77. Figures 3b depicts the ROC under 3 different background level and with 3 replicates per test group, and XBSeq evidently outperformed DESeq as we expected when XBSeq utilized additional non-exonic read count information to estimate the true signal. Further examination under highly expressed (>75% quantile) (Fig. 3c, Supplementary figure S1) and lowly expressed genes (<25% quantile,) (Fig 3d, Supplementary figure S1) condition revealed that among highly expressed genes XBSeq performs only slightly better than DESeq, while XBSeq has much better AUC than DESeq among lowly expressed genes (Supplementary table S3), indicating the importance of background estimation for true signal estimation. We also compared the performance of XBSeq with DESeq (DESeq2), edgeR and edgeR (edgeR_robust), baySeq, limma and EBSeq (Figs 3e and 3f) under high background (high non-exonic read count). As demonstrated in Fig. 3e (3 replicate RNA-seq per test group) and 3f (6 replicate RNA-seq per test group), XBSeq outperformed all the other methods (AUC_XBSeq_ = 0.73, other methods around 0.64, Supplementary Table S4). Overall, XBSeq and DESeq performs comparable at baseline level. When the background noise increases, XBSeq is more robust than DESeq and some other differential expression analysis methods, especially when expression level of the gene is low.

**Table 2.** Area under the curve (AUC) of ROC in different background conditions (low, intermediate or high levels of non-exonic mapped reads) using DESeq and XBSeq, when comparing 3 or 6 replicates in each test group. Overall, XBSeq outperforms DESeq, on average in terms of AUC, by 0.11 and 0.12 for 3 and 6 replicates per group, respectively.

| Replicates<br>per group | DESeq<br>Different background levels |  |  | XBSeq<br>Different background levels |  |  |
| --- | --- | --- | --- | --- | --- | --- |
|  | Low | Intermediate | High | Low | Intermediate | High |
| 3 | 0.69 | 0.66 | 0.64 | 0.82 | 0.78 | 0.72 |
| 6 | 0.76 | 0.73 | 0.70 | 0.89 | 0.86 | 0.80 |

### Control of the false discoveries

We examined the number of false discoveries encountered among the top-ranked genes based on *p* values of different statistical methods. Under baseline level, XBSeq performs comparable with DESeq with similar number of false discoveries (Fig. 4a). Under the scenarios of increased background noise, XBSeq has much less false discoveries compared with DESeq (Fig. 4b). Taking the similar approach, we examined the performance of XBSeq and DESeq among highly expressed (>75% quantile) (Fig. 4c) and lowly expressed genes (<25% quantile) (Fig 4d) with 3 replicates per test group. Even through that DESeq picks up similar number of false discoveries among highly expressed genes compared with XBSeq, it has more false discoveries among lowly expressed genes. Comparisons with some other differential expression methods under high background noise level showed that XBSeq is more robust against false discoveries (Figs. 4e and f). At the pre-selected threshold (*p*-value equals to 0.05), XBSeq has a false discovery around 0.3 which is much less than other statistical methods (around 0.75, Fig. S2). Overall, XBSeq is more robust against false discoveries compared to other statistical methods especially for lowly expressed genes.

**Figure 4.**
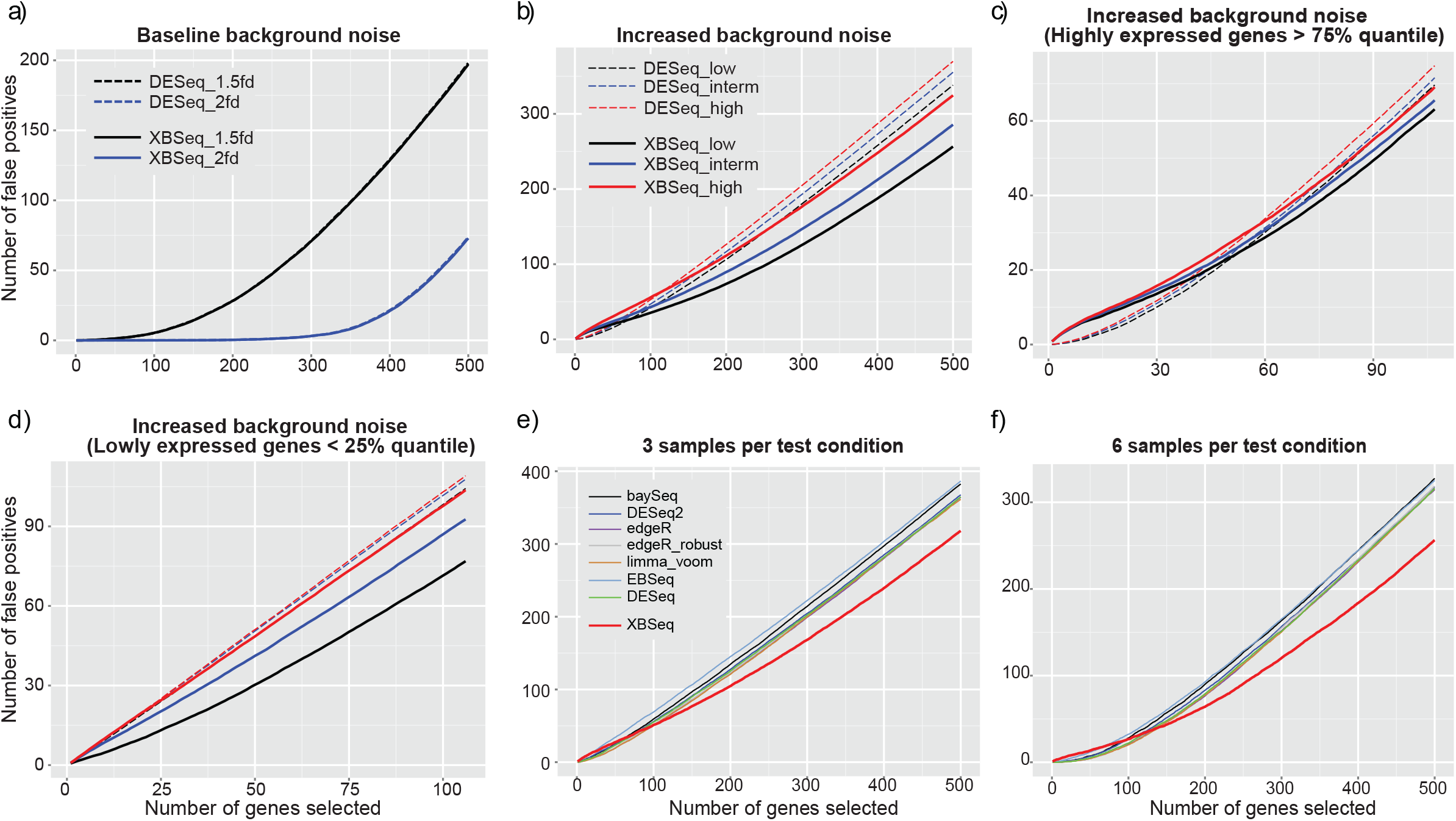
False discovery curves of simulated RNA-seq data in different scenarios. False discovery curves in the scenarios of baseline background noise (a); Different background noises (b); Different background noises with highly expressed genes above 75% quantile of intensity (c); Different background noises with lowly expressed genes below 25% quantile of intensity (d); Comparison with other statistical methods, under high background noise, with either 3 samples per test condition (e), or 6 samples per test condition (f). 3 replicates per test condition, with 10% DEGs and 1.5 fold change were used for (a, b, c, and d).

### Statistical power in detecting DE genes

We compared the performance of XBSeq and DESeq as well as other statistical methods in terms of statistical power at pre-selected threshold (*p* = 0.05) in detecting DE genes (Fig. 5). At baseline level, XBSeq and DESeq perform comparable (statistical power = 0.58 and 0.91 under 1.5 and 2.0 fold change, respectively) (Fig. 5a and Supplementary Tables S1&2). With the increase of the background noise, both methods have decreased statistical power (Fig. 5b and Supplementary Table S3). Even so, XBSeq still has better statistical power at low, intermediate or high background noises. However, different from the result of AUC and false discovery, the statistical power difference between XBSeq and DESeq algorithms among highly expressed genes (>75% quantile, Fig. 5c) and lowly expressed genes (<25% quantile, Fig 5d) are relatively same: both methods perform relatively well (XBSeq is slightly better), and both methods have difficulty in detecting lowly expressed DE genes (with statistical power for XBSeq and DESeq a merely 0.05 and 0.06 respectively, with high background noise). When comparing with some other statistical methods, XBSeq is one of the best methods in terms of statistical power along with algorithms such as edgeR-robust and DESeq2 (Figs. 5e and 5f, Supplementary Table S4). However, with more samples per test group, DESeq2, edgeR and edgeR-robust outperformed XBSeq, due to their robust (or moderate) dispersion estimation. Overall, XBSeq remains one of the best algorithms in terms of statistical power in detecting DE genes compared to other statistical methods.

**Figure 5.**
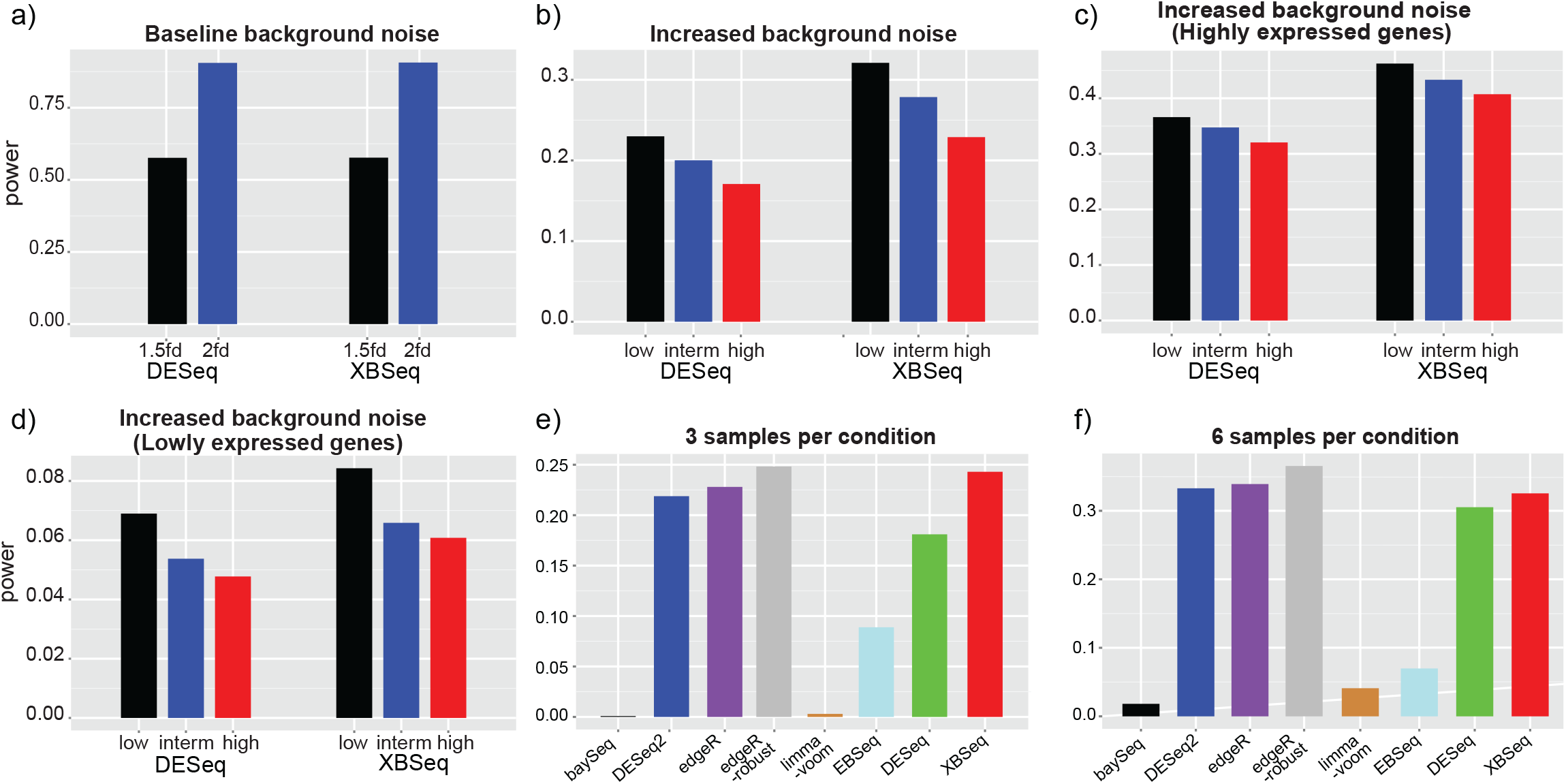
Statistical power of simulated RNA-seq data in different scenarios. Statistical power at pre-selected threshold (*p* = 0.05) in the scenarios of baseline background noise with 1.5 and 2 fold change, with 10% DEGs and 3 replicates per test group (a); Different background noises (b); Different background noises with highly expressed genes above 75% quantile of intensity (c); Different background noises and only lowly expressed genes below 25% quantile of intensity (d); Comparison with other statistical methods, under high background noise, with either 3 samples per test condition (e); or 6 samples per test condition (f). 3 replicates per test condition, with 10% DEGs and 1.5 fold change were used for (a, b, c, and d).

### Application to differential expression analysis for MYC induced gene expression in mouse liver tissues

We have applied a real mouse RNA-seq dataset to different algorithms to test the performance of XBSeq [10]. The mouse RNA-seq dataset includes 3 replicates of wild type mouse (WT) and 3 replicates of Myc transgenic mouse (MYC). Fig. 6 shows a Venn diagram of overlapping and total number of genes detected using XBSeq, DESeq, DESeq2, edgeR, and edgeR_robust with the criterion of *p*-value less than 0.05. XBSeq and DESeq detected similar number of DE genes (446 and 452, respectively, and 414 of them in common, Supplementary Table S5), even if a 1.5 fold change cutoff was added. This is reasonable since the two algorithms share similar differential expression significant test statistics, other than non-exonic read count incorporated into XBSeq. In order to see the different results of XBSeq comparing with others, we listed the exonic and non-exonic read counts in Table 3, which show genes exclusively found by XBSeq (Table 3a) and genes that are not detected by XBSeq but by other algorithms (Table 3b). The venn diagram with 1.5 fold change cutoff added is shown in Fig. S5 and the numbers of overlapped DE genes between XBSeq and other methods are listed in table S5.

**Figure 6.**
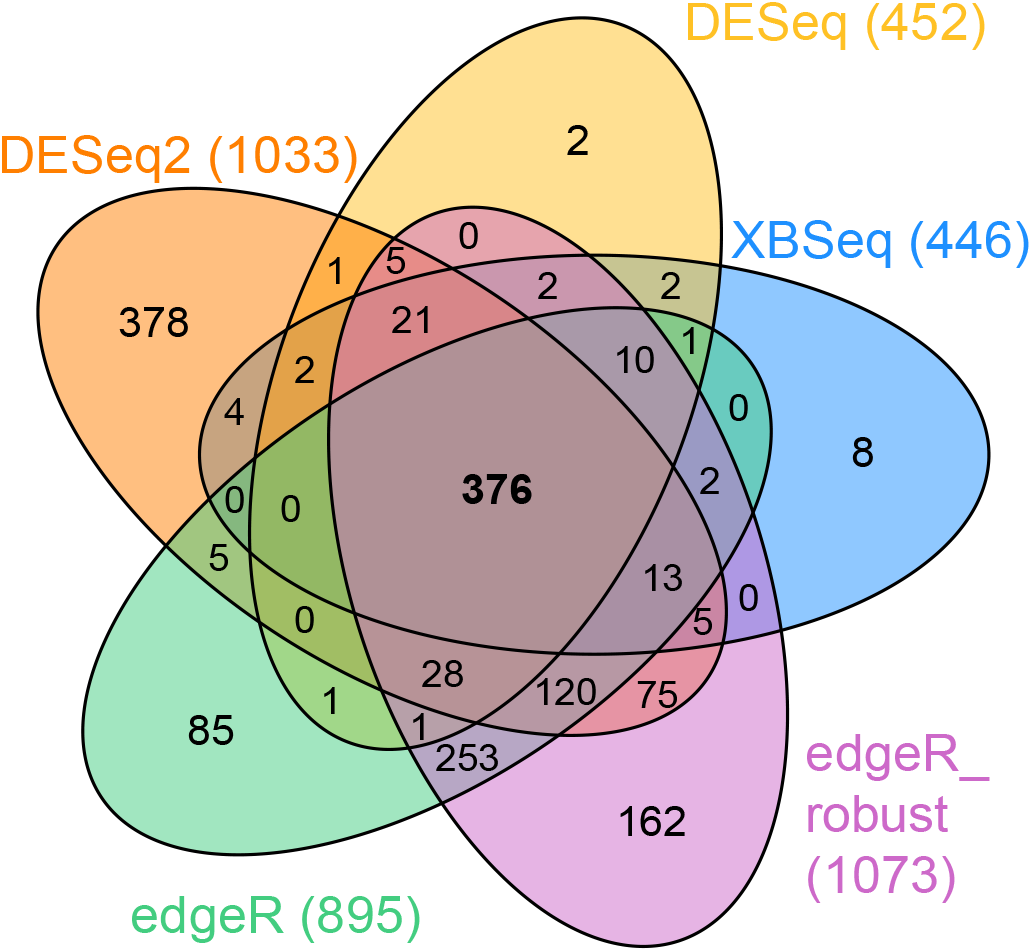
Venn diagram to compare the results of different methods. XBSeq, DESeq, DESeq2, edgeR, and edgeR_robust were applied to a set of mouse RNA-seq data to identify differential genes with *p*-value < 0.05.

**Table 3a.** Two differentially expressed genes (*Nat1*, top table, and *Brca2*, bottom table) that are ONLY detected by XBSeq, showing the exonic, non-exonic, and predicted true signals (estimated by using Eq. 3), from 3 biological replicates for each WT and MYC mouse tissue. 1) FC denotes the fold-change of MYC vs WT, and 2) is the *p*-value from DESeq (for Exonic read only) and XBSeq (for predicted true signal).

| Read counts of gene *Nat1* |  |  |  |  |  |  |  |  |  |  |  |  |
| --- | --- | --- | --- | --- | --- | --- | --- | --- | --- | --- | --- | --- |
|  | WT (3 replicates) |  |  |  |  | MYC (3 replicates) |  |  |  |  | FC<sup>1</sup> | *p*<sup>2</sup> |
| | 1 | 2 | 3 | $\mu$ | $\sigma$ | 1 | 2 | 3 | $\mu$ | $\sigma$ | | |
| Exonic | 63 | 47 | 77 | 62.3 | 15.0 | 91 | 110 | 120 | 107.0 | 14.7 | 1.3 | 0.15 |
| Non-exonic | 20 | 20 | 28 | 22.7 | 4.6 | 23 | 14 | 34 | 23.7 | 10.0 |  |  |
| Predicted true signal | 43 | 27 | 49 | 39.7 | 11.4 | 68 | 96 | 86 | 83.3 | 14.2 | 1.6 | 0.05 |

**Table 3a.** Two differentially expressed genes (*Nat1*, top table, and *Brca2*, bottom table) that are ONLY detected by XBSeq, showing the exonic, non-exonic, and predicted true signals (estimated by using Eq.
| Read counts of gene *Brca2* |  |  |  |  |  |  |  |  |  |  |  |  |
| --- | --- | --- | --- | --- | --- | --- | --- | --- | --- | --- | --- | --- |
|  | WT (3 replicates) |  |  |  |  | MYC (3 replicates) |  |  |  |  | FC<sup>1</sup> | *p*<sup>2</sup> |
| | 1 | 2 | 3 | $\mu$ | $\sigma$ | 1 | 2 | 3 | $\mu$ | $\sigma$ | | |
| Exonic | 213 | 88 | 165 | 155.3 | 63.1 | 132 | 191 | 188 | 170.3 | 33.2 | 0.9 | 0.57 |
| Non-exonic | 216 | 77 | 142 | 145.0 | 69.5 | 90 | 108 | 101 | 99.7 | 9.1 |  |  |
| Predicted true signal | 0 | 11 | 23 | 11.3 | 11.5 | 42 | 83 | 87 | 70.7 | 24.9 | 4.3 | 0 |

**Table 3b.** Two differentially expressed genes (*Calcr*, top table, and *Adh7*, bottom table) that are NOT detected by XBSeq, showing the exonic, non-exonic, and predicted true signals (estimated by using Eq. 3), from 3 biologial replicates for each WT and MYC mouse tissue. 1) FC denotes the fold-change of MYC vs WT, and 2) is the *p*-value from edgeR-robust (for Exonic read only) and XBSeq (for predicted true signal). *Calcr* was detected by all other four methods and *Adh7* was detected by three methods except DESeq (*p*-value = 0.06).

| Read counts of gene *Calcr* |  |  |  |  |  |  |  |  |  |  |  |  |
| --- | --- | --- | --- | --- | --- | --- | --- | --- | --- | --- | --- | --- |
|  | WT (3 replicates) |  |  |  |  | MYC (3 replicates) |  |  |  |  | FC<sup>1</sup> | *p*<sup>2</sup> |
| | 1 | 2 | 3 | $\mu$ | $\sigma$ | 1 | 2 | 3 | $\mu$ | $\sigma$ | | |
| Exonic | 22 | 4 | 20 | 15.3 | 9.9 | 68 | 68 | 64 | 66.7 | 2.3 | 3.3 | 0 |
| Non-exonic | 64 | 4 | 64 | 44.0 | 34.6 | 193 | 218 | 199 | 203.3 | 13.1 |  |  |
| Predicted true signal | 0 | 0 | 0 | 0 | 0 | 0 | 0 | 0 | 0 | 0 | 0 | 1 |

**Table 3b.** Two differentially expressed genes (*Calcr*, top table, and *Adh7*, bottom table) that are NOT detected by XBSseq, showing the exonic, non-exonic, and predicted true signals (estimated by using Eq.
| Read counts of gene *Adh7* |  |  |  |  |  |  |  |  |  |  |  |  |
| --- | --- | --- | --- | --- | --- | --- | --- | --- | --- | --- | --- | --- |
|  | WT (3 replicates) |  |  |  |  | MYC (3 replicates) |  |  |  |  | FC<sup>1</sup> | *p*<sup>2</sup> |
| | 1 | 2 | 3 | $\mu$ | $\sigma$ | 1 | 2 | 3 | $\mu$ | $\sigma$ | | |
| Exonic | 633 | 438 | 429 | 500.0 | 115.3 | 827 | 1149 | 708 | 894.7 | 228.2 | 1.5 | 0.01 |
| Non-exonic | 430 | 241 | 322 | 331.0 | 95 | 643 | 771 | 513 | 642.3 | 129.0 |  |  |
| Predicted true signal | 203 | 197 | 107 | 169.0 | 53.8 | 184 | 378 | 195 | 252.3 | 109.0 | 1.2 | 0.53 |

Specifically, in Table 3a, from only exonic mapped reads, there are no significant difference between WT and MYC samples for genes *Nat1* and *Brca2* (fold change is 1.3, and 0.9 respectively). After subtracting the non-exonic mapped reads, the predicted true signals was significantly differentially expressed between MYC and WT for the two genes (*p*-value = 0.0495, 6×10^−6^ for *Nat1* and *Brca2,* respectively) with fold change increased to 1.6 and 4.3 for *Nat1* and *Brca2*, respectively, along with shrunken standard deviation. From *Brca2*, we can even see that the high dispersion in WT samples are possibly caused by sequencing noises, the gene is barely expressed in WT group as predicted by XBSeq.

On the other hand, with the information in non-exonic regions, XBSeq avoided picking genes that are potentially falsely identified as DE genes because of the background noises. The estimate of the true signal, after considering the non-exonic read counts, may decreased the differential expression and diminish the significance probability. In Table 3b, the two genes, *Calcr* and *Adh7*, were detected by all other four methods, except DESeq for *Adh7* (*p*-value = 0.06), whereas XBSeq tested on the true expression signals and considered them as insignificant changes (*p*-value = 1 and 0.53, respectively).

## Discussion

### Sources of non-exonic mapped reads

Previous studies have shown that non-exonic mapped reads account for about 4~6% of all uniquely mapped reads in mammals [15, 16]. Sequence reads that mapped outside of exonic regions might be originated from different RNA sources depending on the RNA-seq experimental protocol selected for sequencing library preparation. In addition, these non-exonic reads might be derived from experimental artifacts, like genomic DNA contamination, sequencing errors, or unprocessed RNAs, like pre-mRNAs [17], or even non-coding RNAs [18]. van Baker, *et al* [19] has also demonstrated that most of the non-exon mapped reads are associated with the nearby known genes, which suggests that non-exon mapped reads are contextually specific to the corresponding gene. Besides, Hebenstreit, *et al* [20] has shown that all genes from RNA-seq can be classified into two distinct groups, and one of them is the the lower expressed group that consists of putative non-functional mRNAs. All these suggested the biological relevance of incorporating information from non-exonic regions. We have carefully evaluated the aforementioned biological relevance of RNA species in non-exonic regions, and thus necessary steps have been taken for identifying non-exonic regions as discussed in the Methods Section. As we showed in Fig. 1B in one of our real RNA-seq data set, the non-exonic read counts for all genes are mostly less than 0 (log2 RPKM unit), indicating little or no influence from high-expression. We also examined the correlation between the reads mapped to exonic regions and the reads mapped to non-exonic regions in a real RNA-seq experiment. The average correlation is 0.32 which potentially indicates that the non-exonic reads are not ‘functional’ reads which can be used to represent the background noise. By measuring the reads mapped to the non-exonic regions of its corresponding gene but carefully avoid those functional relevant regions, we are able to gain a more reliable estimation of true expression level by eliminating the impact of background noise.

### Compare of XBSeq and DESeq at baseline level

Our simulation at baseline level suggests that XBSeq and DESeq’s performance are virtually indistinguishable in terms of AUC, number of false discoveries and statistical power at baseline level. Not surprisingly, comparison with 6 replicates per test group performed better than 3 replicates per test group even with low level of fold change (fold change at 1.5). Also at baseline background level, the number of truly differentially expressed genes has little effect on the performance of XBSeq and DESeq except with the number of false discoveries. As expected, simulation of 30 percent of true DE genes is more likely to generate false positives than those of 10% true DE genes. However, this effect is dampened with increased number of replicates per test group for differential expression analysis (Table S2).

### Comparison of statistical methods with increased non-exonic mapped reads

To further demonstrate the robustness of XBSeq at different background levels, we simulated genes with low, intermediate or high levels of non-exonic mapped reads. XBSeq outperform DESeq with larger AUC (Figs. 2b-2d), and with better controlling of false discoveries (Figs. 3b-3d). While we achieved excellent true positive detection, we also examined the false negative rate for XBSeq and DESeq, or the statistical power. As shown in Figs. 4b-4d, XBSeq outperform DESeq’s statistical power in detecting DE genes in all three increased background levels. All these suggest that XBSeq is more robust in detecting DE genes in noisy NGS-seq samples.

We also compared performance of XBSeq with some other RNA-seq algorithms, including DESeq2 and edgeR-robust. XBseq excelled in overall performance (better AUC, Figs. 3e & 3f) and false discovery control (Figs. 4e and 4f). XBSeq is also one of the best algorithms in terms of statistical power (Figs. 5e and 5f), indicating a modest trade-off in false negative while maintaining overall performance in DE detection. Moreover, XBSeq performed better with higher non-exonic mapped reads (in all three simulated increased background noise level above baseline). Further examination of algorithms performance among higher (> 75% percentile) or lower expressed genes (< 25% percentile) revealed that false positive genes were mostly generated among lower expressed genes. Among lower expressed genes, XBSeq outperformed than DESeq with better AUC and false discovery. However, both XBSeq and DESeq performed poorly among low-expression genes.

Simulations were carried out with the assumption of NB model for gene expression and Poisson model for background noise (or non-exonic read counts), which is the model XBSeq is built on. While we demonstrated the validity of the assumption in one of our RNA-seq data set (Fig. 1B), we also tested the XBSeq under the uniform or normal model for background read counts while keeping the same NB model for gene expression. As shown in Fig. S2A, for all 3 background distributions (including Poisson model), XBSeq performed better than DESeq, indicating the robustness even when underlying assumption is deviated from Poisson distribution. Another way to examine model bias is by examining the distribution of *p* value under null hypothesis (with no DE genes). As shown in Fig S3, the *p* values generated by most differential expression analysis algorithms are close to uniform distribution from 0 to 1 (showed only 0 to 0.2 in Figs S3), with the exception of EBSeq.

### Execution time of XBSeq algorithm

XBSeq algorithm complexity is similar to DESeq. However, XBSeq will not only take read counts from each gene, but also read counts from non-exonic read counts for each gene, and then perform true signal prediction before evaluating differential expression significance. We benchmarked a set of differential analysis algorithms for their computational times with different number of samples in each condition (Fig. S4). BaySeq algorithm requires the most computational time, followed by DESeq and XBSeq.

## Conclusions

We have developed an approach to take into consideration of non-exonic mapped reads as sequencing noise for precise differential expression analysis. When there is no or very low background noise, the performance of XBSeq is similar to DESeq. However, XBSeq excels when background noise (non-exonic read counts) are higher due to the model estimation of true signal by removing the noise impact. Overall, XBSeq algorithm is shown to be more robust than existing differential expression analysis methods particularly when sequencing noise is a concern.

## Availability of supporting data

The R package of XBSeq, the shift-gene gtf files as well as reproducible scripts for simulation are available from GitHub, https://github.com/Liuy12/XBSeq.

## Authors’ contributions

All authors contribute to the manuscript. HHC, YL, YH and YC conceived and designed the study. HHC and YC implemented the algorithm in Matlab and YL migrated it to R. YL carried out the simulation procedure. YZ designed non-exonic regions for human and mouse, DS contributed biological samples and ZL and YC performed sequencing and basic bioinformatics analysis. All authors read and approved the final manuscript.

## Acknowledgements

This research was supported in part by the Genome Sequencing Facility of the Greehey Children’s Cancer Research Institute, UTHSCSA, which provided RNA-seq service. Fundings for this research were provided partially by the Cancer Prevention and Research Institute of Texas (RP-120685-C2 and RP-120715-C4) YC, National Institutes of Health Cancer Center Shared Resources (NIH-NCI P30CA54174) to YC, NIGMS (R01GM113245) to YH and YC, and the National Science Foundation (CCF-1246073 to YH).

## List of abbreviations

AUC: Area Under the Curve

DE: Differential Expression

NB: Negative Binomial (distribution, model)

NGS: Next Generation Sequencing

ROC: Receiver Operating Characteristic

RPKM: Reads Per Kilobase of transcript per Million reads mapped

## Declaration

The publication costs for this article were funded by the aforementioned CPRIT grants to YC.

